# Network Modules Driving Plant Stress Response, Tolerance and Adaptation: A case study using Abscisic acid Induced Protein-protein Interactome of *Arabidopsis thaliana*

**DOI:** 10.1101/073247

**Authors:** Khader Shameer, Mahantesha Naika, Oommen K. Mathew, Ramanathan Sowdhamini

**Affiliations:** National Centre for Biological Sciences (TIFR), GKVK Campus, Bellary Road, Bangalore 560 065, INDIA; Present address: Icahn Institute of Genetics and Genomic Sciences, New York, NY 10065, USA; Department of Plant Biotechnology, University of Agricultural Sciences, GKVK Campus, Bellary Road, Bangalore 560 065, INDIA

## Abstract

Understanding key protein-protein interaction network mediated by genes responsive to biotic and abiotic stress could help to understand the functional modules and network topologies driven genes responsive to stresses. It still remains to be an open question whether distinct protein-protein interaction networks have functional or regulatory role in mediating abiotic or biotic stress response in plants. To address this question we compiled abscisic acid responsive genes from Stress-responsive TranscrIption Factor DataBase (version 2; STIFDB2); derived protein-protein interaction network mediated by the genes from STRING database and performed biological network analyses using Cytoscape plugins. We have used Molecular Complex Detection algorithm for deriving highly connected module from the abscisic acid responsive network. Biological Network Gene Ontology tool was used to derive functional enrichment of abscisic acid responsive interaction network using GOSlim_Plants ontology. GraphletCounter was used to identify graph motifs in the network and NetworkAnalyzer was used to compute various network topological parameters. We found 26S proteasome subunits as a highly clustered module using Molecular Complex Detection algorithm. Enrichment analysis indicates that several biological processes terms including “flower development” are associated with the network. Results from this case study can be used to understand network properties of abiotic stress responsive genes and gene products in a model plant system.

## Introduction

Plants have prolonged exposure to a variety of abiotic and biotic stresses and such stresses adversely affect the growth of plants. Abiotic and biotic stresses are threat to plants including plant model systems like *Arabidopsis thaliana* and crop species like rice, barley, wheat and corn alike. Stress signals influence the growth of plants including important plant models like *Arabidopsis thaliana* and crop species like rice, barley, wheat and corn (1-8). Several studies in the past have been focused on identifying the role of transcription factors, signaling networks (9,10) and influence of co-expression networks on mediating stress response, tolerance and adaptation in plants (11). Co-expression networks offer an efficient approach to design and interpret network properties, however, due to the limitation in expressing all expressed transcript to proteins due to RNA-modification mechanisms including RNA editing, splicing, and RNA decay pathways. Thus, a canonical protein network could offer a better depiction of network properties (12-15). In the context of ABA-induced stress in *Arabidopsis thaliana*, it still remains to be an open question whether a distinct protein-protein interaction network have functional role in mediating stress response in plants. To address this question we performed biological network analysis of abscisic acid induced stress in *Arabidopsis thaliana* using curated public datasets. Network analysis approaches are currently using in biology to understand collective behavior of molecular entities including genes, proteins and metabolites (16-18).We compiled a dataset of abscisic acid responsive genes from Stress responsive TranscrIption Factor DataBase version 2 (STIFDB2 (19,20)); STIFDB2 have 700 genes upregulated due to ABA from 2 different gene expression studies. We further used the 700 genes as queries and performed a targeted search in STRING database and retrieved all known or predicted protein-protein interaction partners of the 700 genes, where interactant will be one of the 700 genes upregulated due to ABA induced stress. The targeted PPI retrieval approach was used to generate a network with 65 nodes and 108 edges. This network was used as an input to identify clustered modules, network motifs, topological properties and enriched GOSlim_Plants terms.

## Materials and Methods

### Data compilation

Stress responsive genes associated with abscisic acid was compiled from STIFDB2. STIFDB2 is an updated version of a database that catalogued abiotic stress-responsive genes in *Arabidopsis thaliana* (21). The current version of database have a total of 3150 stress responsive genes from *Arabidopsis* transcriptome, 19 stress-responsive transcription factors (ABRE_ABI3_VP1, AuxRE_ARF, C_ABRE_bZIP, DREB_AP2_EREBP, GCC_box_AP2_EREBP, G_ABRE_bZIP, G_box1_bZIP, G_box2_bZIP, G_box_bHLH, HBE_HB, HSE1_HSF, Myb_box1_MYB, Myb_box2_MYB, Myb_box3_MYB, Myb_box4_MYB, Myb_box5_MYB, N_box_bHLH, Nac_box_NAC and W_box_WRKY) and 13 different stress signals (abscisic acid, aluminum, cold, cold-drought-salt, dehydration, drought, high light, iron, NaCl, osmotic stress, oxidative stress, UV-B and wounding). Along with curated stress-responsive genes from public gene expression database Gene Expression Omnibus (GEO) (22), database also have information about predicted transcription factor binding sites in the upstream and 5’ and 3’ untranslated regions (UTR) using hidden-markov model based STIF algorithm (23). The phytohormone abscisic acid (ABA) plays an essential role in adaptive stress responsive pathways and regulates the expression of various abiotic stress-responsive genes (24-31). We compiled a list of 700 genes curated from two different gene expression studies from GEO (GSE23301 (24), GSE9646 (32,33)). List of 700 differentially upregulated due to ABA induced stress can be can be accessed from STIFDB using the URL: http://caps.ncbs.res.in/cgibin/mini/databases/stifdb2/fetch_stress_matrix.pl?se=ABA&orgid=aratha. Detailed aspects of biocuration and genomic data mining steps used to develop STIFDB (21) and STIFDB2 (Mahantesha et. al; accepted in Plant & Cell Physiology) is described elsewhere. The list of 700 genes were queried systematically against a local version of *Arabidopsis thaliana* protein-protein interaction data from STRING (version 9.0) (34) to create an interactome mediated by gene products responsive to ABA induced stress.

### Biological Network Analyses

An interaction network is constructed as follows. For each upregulated gene due to the stress induced by ABA, its first degree interact ants were retrieved from STRING (version 9). If each interactant is upregulated for ABA induced stress – it is retained in the network. Using this targeted protein-protein data retrieval approach an ABA induced stress-responsive network is constructed (see Figure 1 (a)) The resulting protein-protein interaction network was used an input for downstream analyses using Cytoscape (35) and various plug-ins. Network analysis methodology employed in this chapter is summarized in the flow-chart (Figure 1) and brief description tools used in this chapter is provided in Table 1.

**Figure 1:**
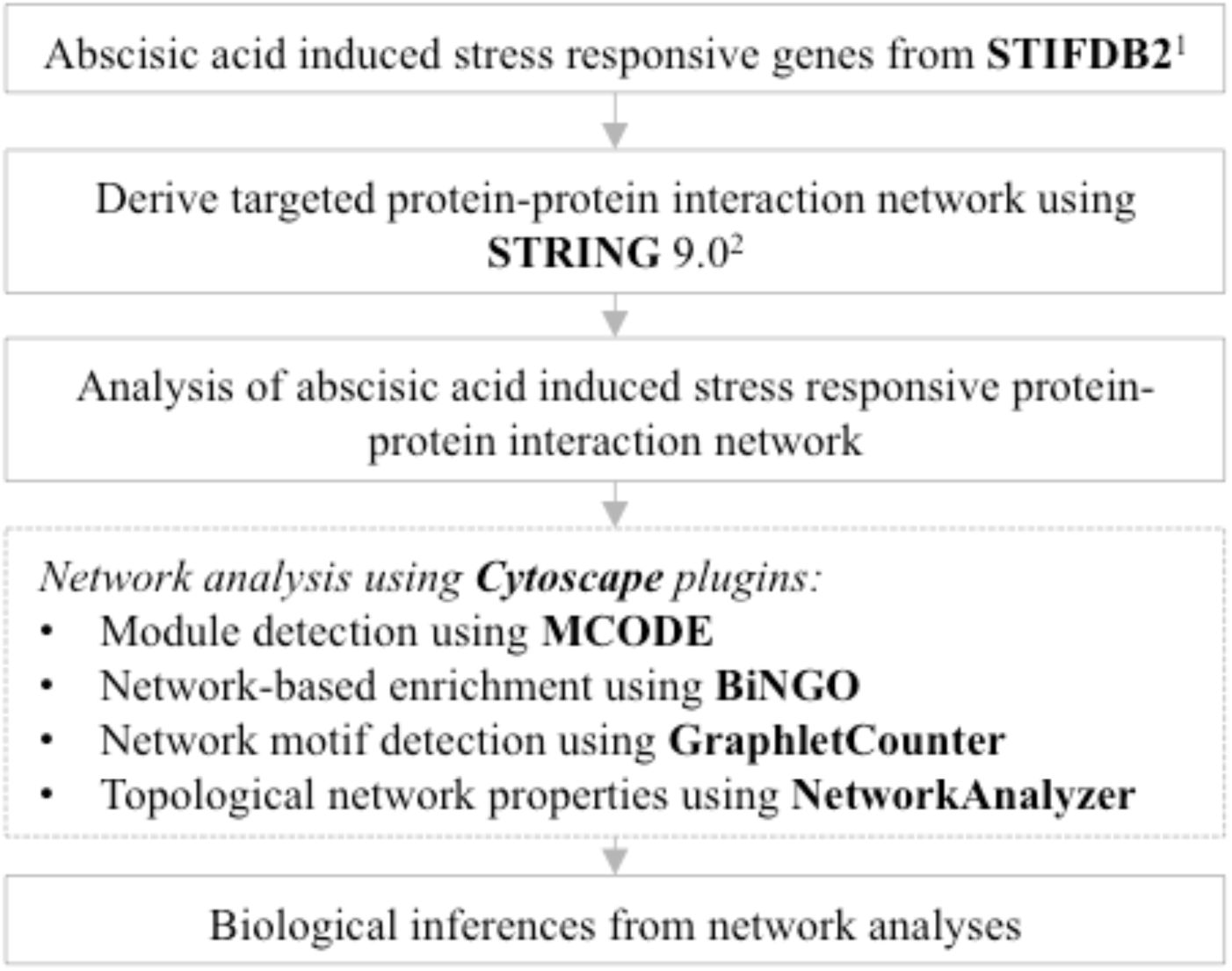
Bioinformatics pipeline used analysis of ABA induced stress responsive protein-protein interaction network. Databases are accessible using URLs: ^1^http://caps.ncbs.res.in/stifdb2; ^2^http://string-db.org

**Table 1:** Brief descriptions of databases and tools used for biological network analysis of ABA induced stress-responsive.

| **Tool / Database** | **Application** |
| --- | --- |
| STIFDB,<br>STIFDB2 | Database of abiotic stress-responsive transcription factors and its putative binding sites on differentially upregulated genes in expression studies |
| STRING | Database of predicted and experimentally validated protein-protein interactions |
| Cytoscape | A software eco-system for analysis and visualization of biological networks |
| MCODE | Cytoscape plugin for identification of clusters, modules or sub-graphs from biological networks |
| GraphletCounter | Cytoscape plugin for identification of graphlets or network motifs |
| BiNGO | Cytoscape plugin for enrichment analysis of components (genes or proteins) in a biological network |

## Identification of core network module mediated by ABA induced stress responsive interactome

A highly interconnected sub-graph embedded in a protein interaction network could suggest the influence of biological principles mediated via protein complex, pathway, coexpression or evolutionary relationships via protein family or superfamily level classifications. We explored the ABA responsive interaction network using molecular complex detection algorithm via the Cytoscape plugin MCODE (36) to identify clustered sub-graphs from the protein-protein interaction network.

## Functional enrichment analysis of ABA induced stress responsive network

Enrichment analysis typically requires a test-set and background set to perform statistical evaluation. In this study the genes associated ABA induced stress responsive network was defined as test-set and background set was defined using annotations available in GOSlim_Plants for *Arabidopsis thaliana.* GOSlim_Plants ontology enrichment analysis was performed using the Cytoscape plugin Biological Network Gene Ontology tool (BiNGO) (37). *P*-values were derived using hypergeometric test, and multiple testing correction was performed using Benjamini and Hochberg false discovery rate (FDR) method. Integrating enriched GO terms based functional enrichment with plant ontology based phenomic enrichment could also provide a distinct view of how genome-phenome elements coordinate to develop stress response, adaptation and tolerance phenotypes (38). Phenomic analyses methods including phenome-wide enrichment analyses have been shown to reveals pleiotropic of genetic variants and genes in human genomes, and these approaches can be applied to plant genomes for better inferences (38-41).

## Network motifs in the ABA responsive network

Similar to sequence and structural motifs in proteins network motifs can be derived from a protein interaction network defined as a graph. Graphlets are defined as a new class of network motifs (sub-graphs with in a large-interactome) for examining local structures within large networks (42). We used GraphletCounter (43,44) to identify the network motifs in the ABA responsive interactome. GraphletCounter enables the identification of graphlets in biological networks. We performed a de-novo motif finding and identified that several graphlets were present in the interactome mediated by ABA responsive genes.

## Topological network properties of ABA responsive network

Quantitative description of the topological properties of network can help to understand the collective nature of ABA responsive network. We computed the following parameters using ABA responsive interactome: clustering coefficient, average number of neighbors, characteristic path length, connected components, isolated nodes, network centralization, network density, network diameter, network heterogeneity, network radius, number of self-loops and shortest paths using the Cytoscape plugin NetworkAnalyzer. Detailed explanation of these parameters can be found elsewhere (45,46).

## Results

Using the targeted PPI retrieval approach was used to generate a network with 65 nodes and 108 edges. This network (Figure 2(a)) was used as query to identify modular complexes (Figures 2(b) and 2(c)) and graphlets (Figure 2(d)). GOSlim_Plants terms enriched among the components of the network was visualized in Figure 3. Using MCODE we identified a cluster with 7 nodes and 15 edges with a score of 2.143. Details about the genes participated in the modular network is provided (Table 2). Our initial analysis indicates that the network is composed of components of 26S proteasome. 26S proteasome was know to play an important role in abiotic stress response including light and oxidative stresses (47-53). Primary function of the 26S proteasome complex is in the clearance of ubiquitinated proteins from the cellular environment for degradation (54). We identified multiple graphlets in the interactome mediated ABA responsive genes. Histogram depicting the distribution of graphlets is provided (see Figure 1(d)). Additional functional or systems biology analyses are required to understand the individual role of proteins associated with the network motifs in abiotic stress response and tolerance. We computed various network topological parameters and compared the properties with a random network derived from *Arabidopsis thaliana* proteome background permuted by keeping the number of edges constant (Table 3). Various network parameters provides insights into topological properties of the stress-responsive network. GOSlim_Plants enrichment analysis results derived from BiNGO is summarized in Table 4 and visualized using Cytoscape (Figure 3).

**Figure 2:**
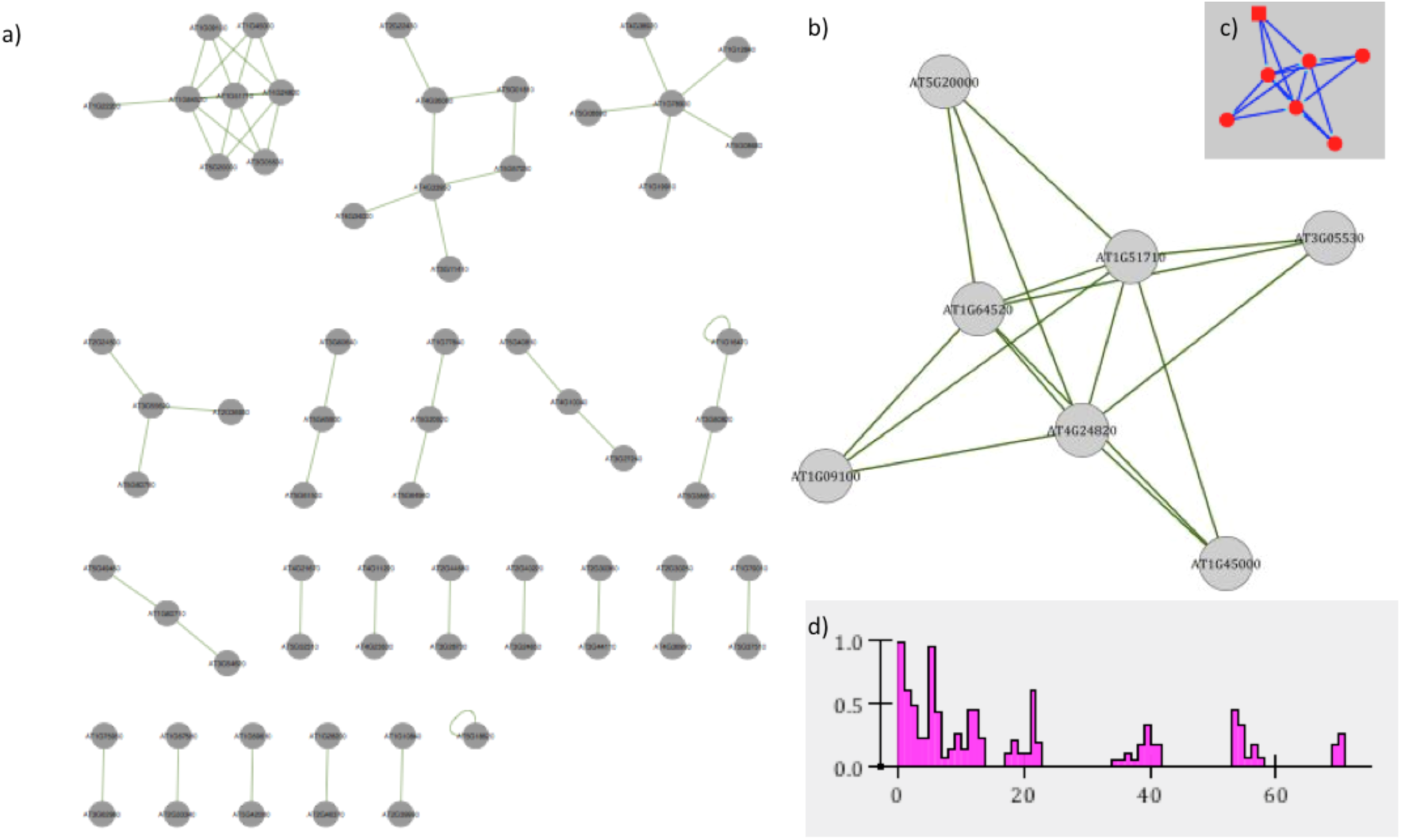
2(a) Protein-protein interaction of ABA induced stress responsive genes derived from STRING. 2(b) Modular protein complex detected using MCODE algorithm visualized using Cytoscape. 2(c) Topology of protein complex detected using MCODE. 2(d) Distribution of graphlets identified using GraphletCounter (*x*-axis indicates the serial number of network motifs used for search, *y*-axis indicates the frequency of individual sub-graphs.

**Figure 3:**
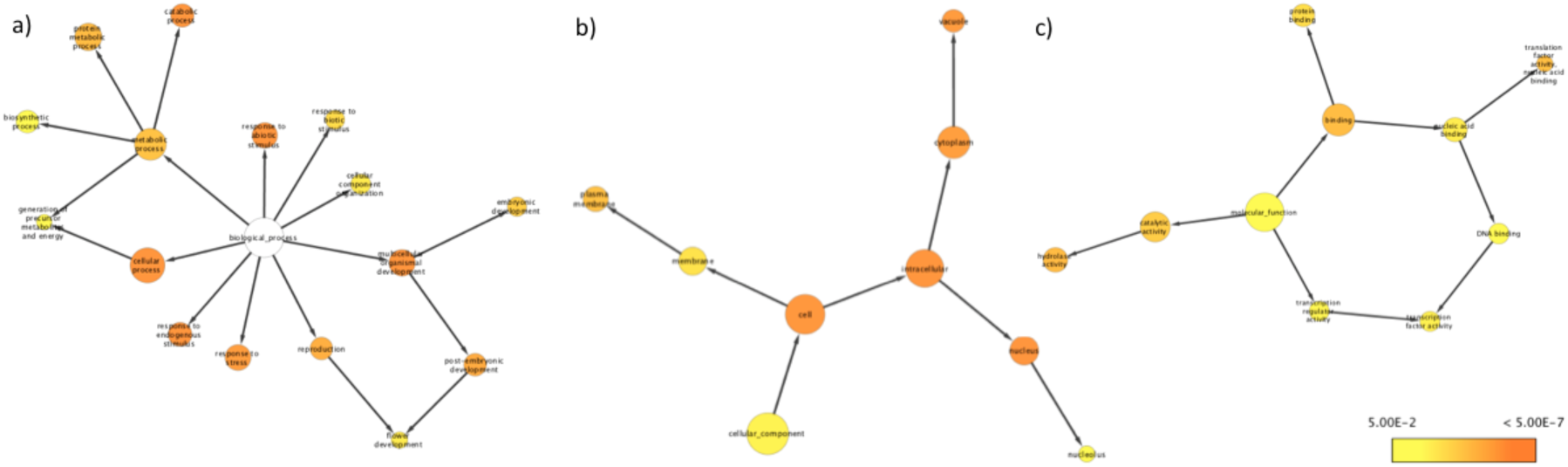
Visualization of GOSlim_Plants terms enriched among the interacting proteins. Heat map indicates *P*-value of individual terms. Nodes in the figure represents terms from 3(a) Biological process, 3(b) cellular components and 3(c) molecular function categories and edge indicates relationship between two terms derived from diacyclic graph of GOSlim_Plants ontology.

**Table 2:** Components of network identified from ABA induced stress responsive protein-protein interaction network.

| **TAIR Locus** | **Chr.** | **Gene name** | **Description** |
| --- | --- | --- | --- |
| AT1G09100 | 1 | RPT5B | 26S proteasome regulatory subunit T5 |
| AT1G45000 | 1 | AT1G45000 | AAA-type ATPase family protein |
| AT1G51710 | 1 | UBP6 | Ubiquitin carboxyl-terminal hydrolase 6 |
| AT1G64520 | 1 | RPN12a | 26S proteasome non-ATPase regulatory subunit RPN12A |
| AT3G05530 | 3 | RPT5A | Regulatory particle triple-A ATPase 5A |
| AT4G24820 | 4 | AT4G24820 | 26S proteasome regulatory subunit N7 |
| AT5G20000 | 5 | AT5G20000 | AAA-type ATPase family protein |

**Table 3:** Topological properties of ABA responsive interactome derived using NetworkAnalyzer.

| **Network parameters** | **Results from ABA network** | **Random network<sup>#</sup>** |
| --- | --- | --- |
| Clustering coefficient | 0.087 | 0.0 |
| Connected components | 22 | 98 |
| Network diameter | 3 | 3 |
| Network radius | 1 | 1 |
| Network centralization | 0.087 | 0.010 |
| Shortest paths | 194(4%) | 240(0%) |
| Characteristic path length | 1.505 | 1.108 |
| Avg. number of neighbors | 1.631 | 1.049 |
| Number of Nodes | 65 | 206 |
| Network density | 0.025 | 0.005 |
| Network heterogeneity | 0.823 | 0.225 |
| Isolated nodes | 1 | 0 |
| Number of self-loops | 2 | 0 |

<sup>#</sup>Permuted different number of genes from Arabidopsis thaliana proteome by keeping the number of edges constant.

**Table 4:** Enriched GOSlim_Plants terms identified using BiNGO.

| **GO-ID** | **P-value<sup>#</sup>** | **Description** |
| --- | --- | --- |
| GO:5622 | 1.96E-10 | Intracellular |
| GO:9987 | 2.78E-08 | Cellular process |
| GO:9628 | 2.78E-08 | Response to abiotic stimulus |
| GO:9056 | 7.09E-08 | Catabolic process |
| GO:7275 | 7.09E-08 | Multicellular organismal development |
| GO:9719 | 8.96E-08 | Response to endogenous stimulus |
| GO:5634 | 3.73E-07 | Nucleus |
| GO:5623 | 6.11E-07 | Cell |
| GO:5773 | 9.04E-07 | Vacuole |
| GO:6950 | 1.24E-06 | Response to stress |
| GO:5737 | 1.59E-06 | Cytoplasm |
| GO:9791 | 5.64E-06 | Post-embryonic development |
| GO:3 | 8.90E-06 | Reproduction |
| GO:19538 | 3.05E-05 | Protein metabolic process |
| GO:16787 | 3.05E-05 | Hydrolase activity |
| GO:5488 | 3.37E-05 | Binding |
| GO:8135 | 4.98E-05 | Translation factor activity, nucleic acid binding |
| GO:5886 | 5.74E-05 | Plasma membrane |
| GO:8152 | 6.56E-05 | Metabolic process |
| GO:3824 | 1.87E-04 | Catalytic activity |
| GO:9790 | 2.31E-04 | Embryonic development |
| GO:9607 | 1.01E-03 | Response to biotic stimulus |
| GO:5515 | 1.24E-03 | Protein binding |
| GO:16020 | 2.10E-03 | Membrane |
| GO:16043 | 4.39E-03 | Cellular component organization |
| GO:9908 | 5.69E-03 | Flower development |
| GO:3676 | 6.36E-03 | Nucleic acid binding |
| GO:3700 | 8.05E-03 | Transcription factor activity |
| GO:5575 | 1.24E-02 | Cellular component |
| GO:30528 | 1.81E-02 | Transcription regulator activity |
| GO:6091 | 2.65E-02 | Generation of precursor metabolites and energy |
| GO:3677 | 2.96E-02 | DNA binding |
| GO:5730 | 3.06E-02 | Nucleolus |
| GO:3674 | 3.42E-02 | Molecular function |
| GO:9058 | 3.58E-02 | Biosynthetic process |

<sup>#</sup> Bonferroni corrected

## Discussion

Several studies in the past have been focused on identifying the impact of transcription factors and influence of co-expression networks on mediating stress response, tolerance and adaptation in plants. Biological network analysis is an emerging sub-discipline of computational systems biology that enable identification of functionally connected modules among genes, proteins, metabolites or other small molecules identified from a primary experiment. In general biological interactions are mediated by macromolecular interactions (protein-protein, protein-DNA and protein-ligand interactions). Protein-protein interaction can be characterized experimentally by performing targeted or high-throughput yeast-two-hybrid screening or protein fragment complementation assays. With the advent of molecular databases capturing interaction data, the data captured from individual assays and high throughput studies can be extrapolated to new genes or proteins in a predictive manner. In this chapter, we illustrated the application of biological network analysis approaches for understanding functional modules mediated by upregulated genes due ABA induced stress. ABA induce a variety of abiotic stresses like oxidative and osmotic stresses and also play a crucial role in biotic and abiotic stress response, tolerance and adaptation. As we are in an era, where detailed experimental characterization of protein-protein interaction due to the abiotic stresses are not available in large number, such analytical approaches could valuable resource to design experiments. The interactome derived from stress responsive genes from the curated database STIFDB2 could be helpful to characterize the complex circuitry with which stress-responsive transcription factors might regulate each other and other proteins during the process of stress response and tolerance. Expanding this analysis to entire set of abiotic and biotic stresses compiled in STIFDB2 for *Arabidopsis* and rice could help to understand various network properties of stress-responsive networks and its role in stress tolerance and adaptation.

## Future directions

In this chapter we explained a strategy to leverage publicly available datasets and curated information compiled in STIFDB2 to study abiotic stress response in plants. We envisage performing this study for multiple stresses in Arabidopsis and other crop species for a better understanding of biological network structure associated with abiotic stress response and stress tolerance events in plants.

## Acknowledgements

The authors thank NCBS, TIFR and UAS (B) for infrastructural support.

